# Spatiochemical Investigation of Potential Biocontrol Agents Against *Phytophthora capsici* Infection in Tomato

**DOI:** 10.1101/2024.12.02.626494

**Authors:** Jamille Y. Robinson, Hawkins S. Shepard, Daniel Ambachew, Peter J. Eyegheleme, Jody C. May, Margaret T. Mmbaga, John A. McLean

## Abstract

Biological control agents can offer an eco-friendly and more sustainable alternative to conventional chemical pesticides, providing protection against destructive pathogens, such as *Phytophthora capsici,* while reducing potential environmental harm associated with synthetic pesticide use in agricultural production systems. This work demonstrates the biocontrol effectiveness of various *Bacillus* species, including *Bacillus vallismortis*, *Bacillus amyloliquefaciens, Bacillus thuringiensis,* and *Bacillus subtilis*, against the widespread plant pathogen, *Phytophthora capsici*. Our studies showed that *Bacillus thuringiensis* and *Bacillus subtilis* promote plant growth and provide protection against *Phytophthora capsici* in both *in vitro* and *in vivo* greenhouse studies, while *Bacillus vallismortis* and *Bacillus amyloliquefaciens* were effective *in vitro* but not *in vivo*. Specifically, *Bacillus thuringiensis* was observed to both hinder the growth of *Phytophthora capsici* and enhance plant resilience to this oomycete pathogen. To probe the molecular interactions between biocontrol agent and pathogen, a dual culture of *Bacillus thuringiensis* and *Phytophthora capsici* was analyzed *in situ* using a mass spectrometry imaging workflow that implemented desorption electrospray ionization. This imaging approach spatially investigated the complex biochemical interactions that serve as the molecular foundation for the effectiveness of these biological control agents in crop protection, including antagonistic interactions that may be fundamental to their method of action. Herein, we demonstrate the benefits of biological control agent application in tomato cultivation, including enhanced pathogen control and plant growth, and showcase the strengths of desorption electrospray ionization-mass spectrometry imaging when applied to the spatially-resolved molecular characterization of agriculturally relevant microorganisms.

## Introduction

*Phytophthora capsici* is a notorious filamentous oomycete plant pathogen that causes significant harm in tomato production, primarily through root rot, fruit rot, and foliage blight. *Phytophthora spp*. primarily release effector proteins through specialized structures called haustoria that are used for infection establishment and absorbing nutrients from the host plant. The effector proteins hamper the plant’s ability to absorb water and nutrients, leading to stunted growth and vigor (Wang et al. 2013; Wang et al. 2018). Considering that *Phytophthora spp*. are the causative agents responsible for some of the most destructive plant diseases, their notoriety makes these organisms excellent models for studying the complex molecular interactions between plants and invading pathogens.

Historically, chemical pesticides have been widely used to control oomycete plant pathogens. However, synthetic pesticides have several shortcomings, such as soil contamination attributed to accumulating chemical residues. These chemicals disrupt the soil microbiome by reducing microbial diversity and altering community structures, impairing essential soil functions. Furthermore, the selective pressure from pesticides fosters the development of resistant pathogen strains, exacerbating pest control challenges and potentially increasing pathogen virulence (Sehrawat et al. 2021; Pathak et al. 2022). Together, these effects contribute to a cycle of declining soil health, increased dependency on chemical inputs, and escalating environmental and agricultural issues. One promising alternative is the use of beneficial bacteria acting as biocontrol agents (BCA), which offer several advantages over traditional chemical pesticides. Specifically, rhizobia-dwelling, plant growth-promoting bacteria (PGPB) protect plants from diseases through a combination of competitive exclusion, production of antimicrobial compounds, enhancement of plant immune responses, and beneficial modifications of the rhizosphere environment (Ahemad et al. 2014; Glick 2012). These mechanisms not only help in controlling pathogenic microbes but also contribute to overall plant health and productivity.

Many PGPBs produce antimicrobial substances such as antibiotics, siderophores, and lytic enzymes that inhibit or kill pathogenic microbes (Narayanan and Glick, 2022). These compounds can directly suppress or kill pathogens, reducing their ability to infect the plant. PGPBs can also trigger a plant’s defense mechanisms through induced systemic resistance (ISR). When plants detect specific signals from PGPB, they activate a wide range of defense responses that enhance their ability to fend off pathogens. These responses include the production of defensive chemicals, strengthening cell walls, and activating stress-related proteins (Kaur et al. 2022).

Additionally, PGPB can prime the plant immune system, making it more responsive to pathogen attacks. This priming induces a heightened state of alertness that allows the plant to respond more quickly and robustly when pathogens attempt to invade. Another protection facet is the ability of PGPB to preemptively occupy and colonize plant root surfaces and internal tissues. This ability can prevent pathogenic microbes from establishing themselves by forming a physical barrier that reduces the chances of infection by pathogens. The PGPB can also alter the rhizosphere’s chemical and physical properties in ways unfavorable to pathogens by modifying pH levels, producing volatile organic compounds, or altering the availability of certain nutrients, making the environment less conducive to pathogen survival and growth (Kaur et al. 2022; Compant et al. 2010).

Bacillus species play a significant role in the biocontrol of plant pathogens through various mechanisms. These beneficial bacteria are known for producing a wide range of bioactive compounds and adapting to different environmental conditions (Noufal et al. 2018; Al-Obaidy 2014; Liu et al. 2015). *In vitro* studies examined the biocontrol effectiveness of *Bacillus spp.* [*B. vallismortis* (Ps), *B. amyloliquefaciens* (Psl), and *B. thuringiensis* (IMC8)] against *P. capsici* in the management of tomato diseases. The results indicated a reduction in mycelial growth attributed to *B. amyloliquefaciens* and *B. vallismortis,* while *in vivo* investigations demonstrated a greater reduction in disease severity compared to the systemic fungicide metalaxyl (Bhusal and Mmbaga, 2020). Biocontrol microorganisms are also credited for antibiosis characteristics that induce plant responses to optimize microbe-based biocontrol methods (Köhl et al. 2019). Strategies that determine the ecological functions of bacterial secondary metabolites, the variables that regulate their production, and their physicochemical properties are all still in their infancy. Substituting synthetic fungicides with bacterial BCAs in plant pathogen management is a viable and auspicious strategy, owing to its ecological impacts, diminished residues on agricultural commodities and the environment, and adaptability across diverse agricultural systems (Rotich et al. 2019; Joshua and Mmbaga 2020). However, before these alternatives can be implemented, further investigation and analysis of the potential mechanism of action is warranted to determine more effective ways to elucidate the metabolome and underlying biochemical processes of BCA-pathogen interactions.

Mass spectrometry (MS) based workflows are frequently implemented in biochemical profiling due to the ability to leverage the high sensitivity, specificity, and speed of MS towards the study of small molecules via metabolomic and lipidomic analyses (Han and Gross, 2014). Increasing attention is being given to mass spectrometry imaging (MSI) approaches, as the spatiochemical information gathered during the imaging process can be used to create molecular maps that inventory the compounds present at every spatial location sampled. These techniques go beyond mere detection and provide important insight into the distribution of specific analytes found within the samples. MSI-based workflows are increasingly utilized in medical fields, imaging tissue sections to characterize disease states, identifying the localization of exogenous compounds, and even providing quantitative data for the concentrations of biomolecules (Gessel et al. 2014; Li et al. 2016; Zhang et al. 2011; Manicke et al. 2009). Among the different implementations of MSI, desorption electrospray ionization (DESI) shows particular promise as it combines the benefits of traditional spray ionization methods with the analysis of condensed phase samples usually associated with desorption-based approaches (Morato et al. 2023). Importantly, DESI experiments can be conducted under ambient conditions, allowing for the direct examination of living biological systems while minimally affecting the analysis of the small molecules required for effective characterization (Ifa et al. 2007).

This study aims to investigate the feasibility of using DESI to examine agricultural biological systems involving biological control of plant pathogens such as *P*. *capsici* using bacterial endophytes such as *B. thuringiensis*. In this study, we evaluated biological agents that can be used to manage *P*. *capsici* and provide environmentally friendly products as alternatives to chemical pesticides. The co-culture of *B. thuringiensis* and *P. capsici* is used to determine if DESI-MSI could serve as a potential analytical strategy for spatial analysis of BCA-pathogen interactions and communication. We compared the effect of specific BCAs on *P. capsici* in tomato plants and found that *B. thuringiensis* exhibited the greatest effects in reducing disease severity and improving plant growth, prompting further characterization of potential mechanisms of action by co-culturing the two organisms. DESI MSI allowed us to directly examine the living co-culture grown on a membrane scaffold *in situ* and provide spatially resolved chemical profiles relating to this interspecies interaction. This study combined the traditional agricultural characterization methodologies with DESI-MSI, and findings reported herein focused on specific BCA interactions between *B. thuringiensis* and *P. capsici* thus providing a potential workflow that can be used for the molecular characterization of agricultural biological control systems.

## Results

### BCA-treated plants exhibit growth promotion and pathogen protection

Two BCA isolates, *B. thuringiensis* (IMC8) and *B. subtilis* (Prt), stood out for growth promotion of tomato seedlings three weeks after BCA treatment (**Fig. 1B, C**). Evaluation of the effect of BCAs on tomato plants in the presence of *P. capsici* pathogen used 10^4^ zoospores per mL and controls not inoculated with *P. capsici*. Plants were moved to the greenhouse environment and the study was terminated 10 weeks after BCA treatments. The BCA-treated plants exhibited similar growth characteristics when exposed to *P. capsici* with biomasses similar to control plants not inoculated with *P. capsici* and heavier than *P. capsici*-treated tomatoes (**Fig. 1D**). Non-BCA-treated plants exposed to *P. capsici* exhibited a considerable decrease in biomass marked by increased infection and only 38% of plants survived after ten weeks. Meanwhile, BCA-treated plants exposed to *P. capsici* had a 75% survival rate for IMC8 and Prt (**Fig. 1E**). Plants treated with *B. amyloliquefaciens* (Psl) and *B. vallismortis* (PS) in the presence of *P. capsici* had a 38% and 25% survival rate, respectively. This indicated that IMC8 and Prt BCAs provided superior plant protection against *P. capsici* compared to PS and Psl. PS-treated plants demonstrated an 88% survival rate. Notably, plants treated with IMC8, Prt, and Psl with no pathogen exposure displayed 100% survival after ten weeks, suggesting that BCA treatment could be offering additional protection from other potential pathogens present in non-sterile soil used in the greenhouse, considering that only 50% of non-BCA-treated control plants remained viable after 10 weeks (**Fig. 1E**).

**Figure 1.**
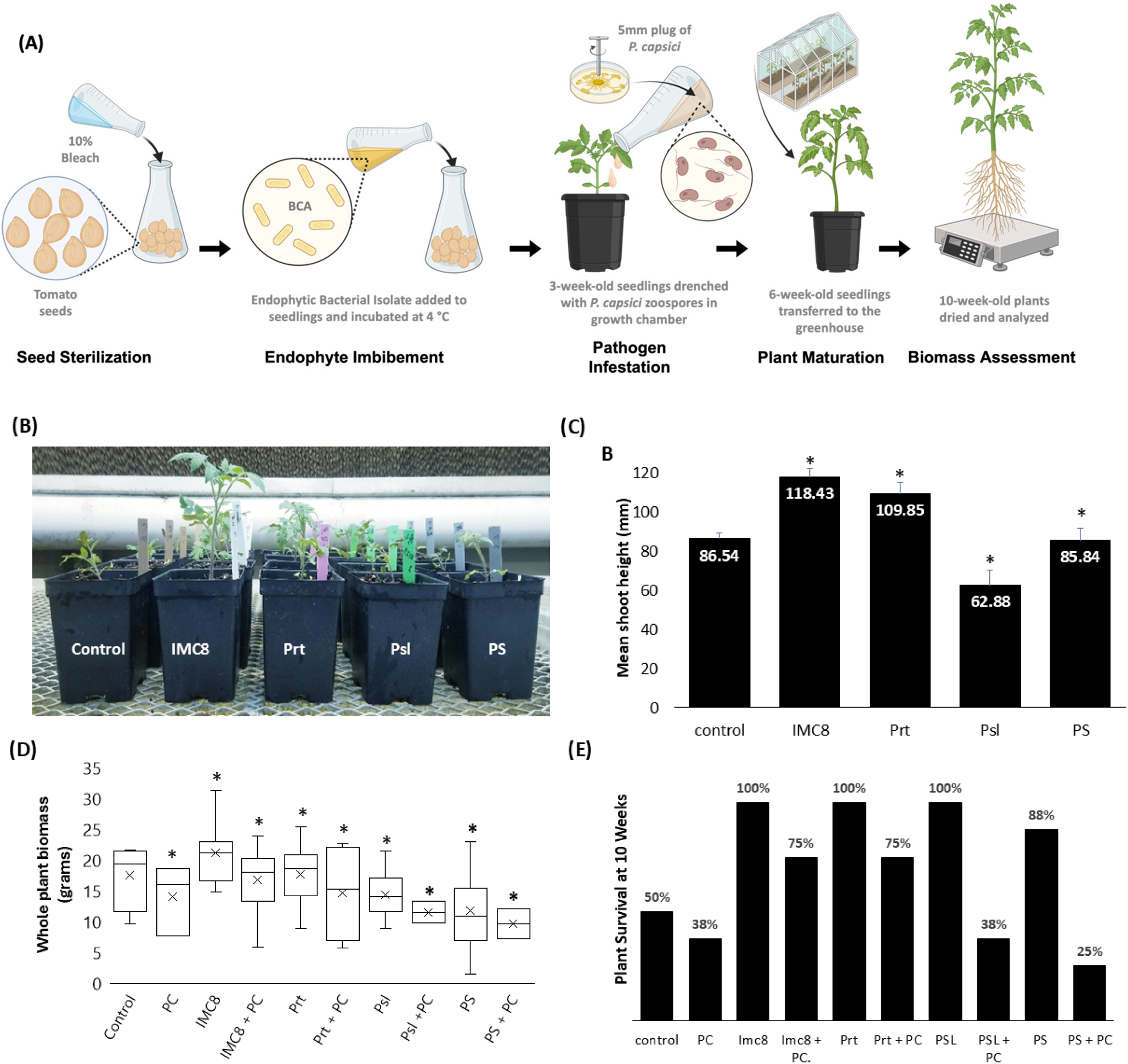
Efficacy of Endophytic biological control agents (BCA) on tomato seedlings inoculated with *Phytophthora capsici*. **(A)** Schematic representation of tomato seed treatment with bacterial isolates using inoculum of 10^4^ CFU/mL. Tomato seeds were sterilized in 10% sodium hypochlorite (Clorox bleach) for 10 min, rinsed three times in sterile water, and blotted dry using heat sterilized paper towels, then soaked in bacterial suspensions of 10^8^ colony forming units (CFU) for 24 hrs at 4 °C. The seeds were then sown in heat-sterilized soil in a growth chamber maintained at 26 ± 3 °C. **(B, C)** Shoots of three-week-old tomato seedlings grown in treated with *Bacillus thuringiensis* (IMC8), *Bacillus subtilis* (Prt), *Bacillus amyloliquefaciens* (Psl), and *Bacillus vallismortis* (PS) with a replication of ten to twelve individual plants per bacterial treatment were measured from root collar (caliper) to terminal bud (meristem); values are means ± SE. *, p-value of <0.05 when compared to BCA and control. **(D)** BCA effect on plant dry biomass of 10-week-old tomato plants inoculated with 10^5^ zoospores/mL *P. capsici* and transferred from growth chamber and transplanted into non-sterile soil in greenhouse environment;. *, p-value of <0.05 when compared between all groups to control. **(E)** The percentage of plants that survived from *P. capsici* root infections in 10-week-old tomato seedlings in greenhouse non-sterilized soil inoculated with 10^5^ zoospores per mL with controls not inoculated with BCA or *P. capsici*.

However, *in vitro* studies revealed that all BCA-treated seedlings exhibited pathogen protection and growth promotion (**Fig. 2**). This observation suggests that IMC8 promoted the plant’s immune system and aided plant protection against *P. capsici,* with the overall plant weight being significantly heavier than the control not treated with BCAs. Bacillus isolates Prt, IMC8, and Psl all have statistically significant (p-value <0.05, 95% confidence interval) growth-promoting effects on tomato shoots *in vitro* (**Fig. 2C**) compared to control after two weeks, but at ten weeks of greenhouse study in the presence of the pathogen, IMC8 exhibited the most biomass increase amongst BCA-treated tomato seedlings in comparison to the nontreated control (**Fig. 1D**). This suggested that growth promotion and disease protection may depend on factors such as BCA and pathogen inoculum concentrations following the initial treatment.

**Figure 2.**
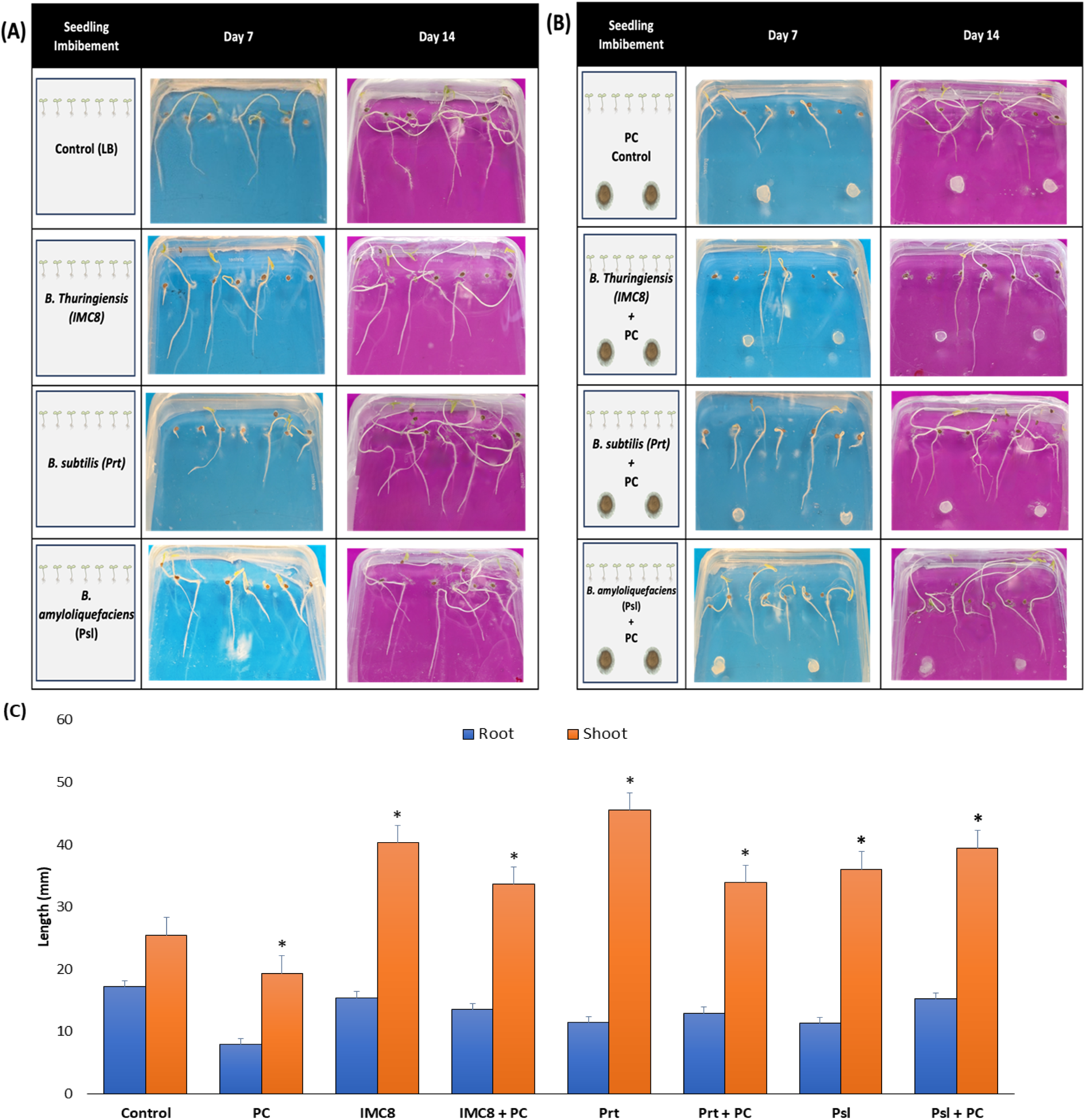
Alterations in the growth of tomato seedlings treated with endophytic biological control agents (BCA) and challenged by *Phytophthora capsici* (PC). Tomato seeds were sterilized in 10% Clorox bleach for 10 minutes, followed by three rinses in sterile water, immersion in bacterial suspensions of 10^8^ colony-forming units (CFU), storage at 4 °C for 24 hours, and subsequent sowing in Murashige & Skoog (MS) growth media agar plates devoid of sugar. Seedlings were maintained in a dark environment at ambient temperatures of 23 °C for 48 hours prior to exposure to light in a growth chamber regulated at 26 ± 3 °C. **(A)** Treatments with *Bacillus thuringiensis* (IMC8), *B. subtilis* (Prt), *B. amyloliquefaciens* (Psl), and *B. vallismortis* (PS) with a replication of seven individual plants per treatment. **(B)** Two 5 mm plugs of actively growing *P. capsici* were placed equal distances apart approximately 35 mm from the bottom of the plate. **(C)** Root and shoot measurements were obtained on days 7 and 14 using Image J; values are means ± SE. *, p-value of <0.05 when compared between all treated groups to control.

### BCA provides immunity prior to pathogen exposure

Tomato seedlings treated with BCAs and exposed to *P. capsici* exhibited longer roots and shoots after 14 days (**Figs. 2A-B**). The lengths of the roots were statistically similar to the nontreated control not exposed to the pathogen or BCA (**Fig. 2C**). However, shoot height more than doubled in plants treated with Prt (*B. subtilis*; 138.6% increase) and IMC8 (*B. thuringiensis*; 110.5% increase). Plants treated with Psl displayed an increase of only 89.5% compared to the nontreated controls. Seedlings exposed to *P. capsici* without BCA treatment displayed a 94.6% decrease in root length and a 24.1% decrease in shoot height. However, IMC8-treated plants exposed to *P. capsici* displayed only a 12.2% decrease in root length. Overall, there was a slight increase in root length of seedlings treated with Prt and Psl and exposed to the pathogen by 13.2% and 34.5%, respectively, however this increase was not statistically significant (p-value <0.05, 95% confidence interval). BCA-treated plants exposed to *P. capsici* pathogen exhibited a significant increase in shoot height by 74%, 75.2%, and 103.9% from IMC8, Prt, and Psl, respectively, compared to the non-BCA-treated plants exposed to *P. capsici*.

### Pathogen mycelium exhibits hyphae collapse in the presence of BCA

Microscopic analysis revealed notable changes in the morphology of *P. capsici* hyphae within the zone of inhibition in response to the BCA after 48 hours of co-cultivation. Control *P. capsici* grown in the absence of BCAs exhibit hyphae that appeared healthy, with smooth, robust, and uniformly tubular structures (**Fig. S1**). By contrast, as *P. capsici* hyphae grew closer to active culture of IMC8, it began to display signs of structural integrity loss, including decreased hyphal branching **(Fig. 3A-D)**. Most prominently, there was widespread hyphal collapse within the zone of inhibition, characterized by the flattening and fragmentation of hyphae (**Fig. 3C-D**). The affected hyphae experienced contraction, and their overall structural stability was reduced, suggesting disruption to the cell wall and membrane. Microscopic examination also showed that the hyphae near *B. thuringiensis* displayed cytoplasmic leakage and vacuolization **(Fig 3D).** These symptoms suggest that there is damage to the cells and a decrease in the internal pressure within the hyphal cells, causing them to collapse. These visual differences observed between the oomycete hyphae suggest that different metabolic states were induced within *P. capsici* when grown under *B. thuringiensis* exposure compared to the control. Variations in the assortment and quantity of small molecules present within a system can typically be correlated to the underlying biology, making the investigation of these small molecules integral for understanding any potentially altered metabolism. As such, the DESI-MSI workflow outlined in **Fig 3E** was applied to gather global metabolite information for both species through a single untargeted acquisition.

**Figure 3.**
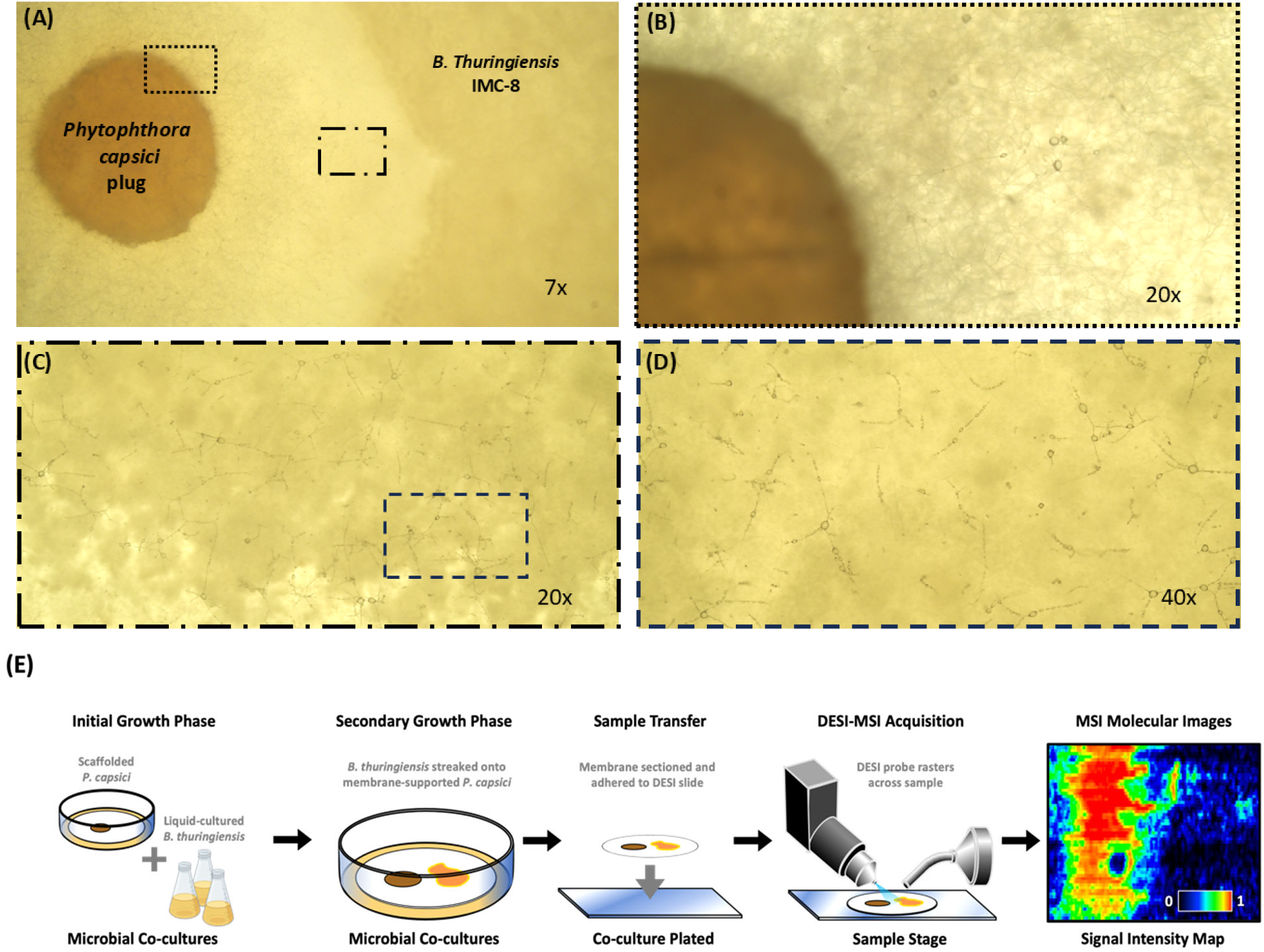
Biological Control Agent (BCA) effect on *Phytophthora capsici* hyphae morphology under brightfield microscopy. **(A)** Bright-field micrographs (7x) of a dual culture containing a 5 mm plug *Phytophthora capsici* growing concomitantly alongside endophyte, IMC8 *(B. thuringiensis*). **(B)** Normal hyphae morphology of *Phytophthora capsici* at 20x. **(C, D)** Zone of inhibition hyphae collapse at 20x and 40x. **(E)** Workflow diagram outlining the sample handling process for the *P. capsici* and *B. thuringiensis* co-culture. Initial culturing of microbes occurs independently, prior to the microporous membrane scaffolding procedure. After co-cultures are grown to the desired point, the membrane is removed from agar, cut to size, and affixed to a glass microscope slide for desorption electrospray ionization mass spectrometry imaging (DESI-MSI) acquisition.

### DESI-MSI molecular characterization and unsupervised segmentation

The spatial distributions of *ca.* 4000 total analyte features were measured for the bacteria-oomycete co-culture during the DESI-MSI acquisition and processed into location-resolved chemical profiles (**Fig. 4B**). In order to identify the spatially relevant and statistically significant features, complete chemical signature information was analyzed using an unsupervised segmentation algorithm (Bemis et al. 2016). This spatially shrunken centroids (SSC)-based k-means clustering algorithm demarcated seven chemically distinct regions within the co-culture, disregarding signals corresponding to noise and analytes with homogenous spatial distributions that are characteristic of chemical background (**Fig. 4B**). By comparing the output of the segmentation algorithm to the optical image of the sample taken prior to imaging (**Fig. 4A**), these seven segments can be qualitatively assigned into three general regions as follows: (1) three segments associated with different phases of BCA growth, (2) three segments of interspecies interaction, and (3) one segment associated with oomycete growth (**Fig. 4D**). The region of interspecies interaction (Segments 2, 3, and 4), is particularly relevant for understanding the altered metabolic state arising in co-culture conditions, as these are the areas in which the two species are in direct contact with one another and correspond to the zone of inhibition identified by microscopy (**Fig. 3A**).

**Figure 4.**
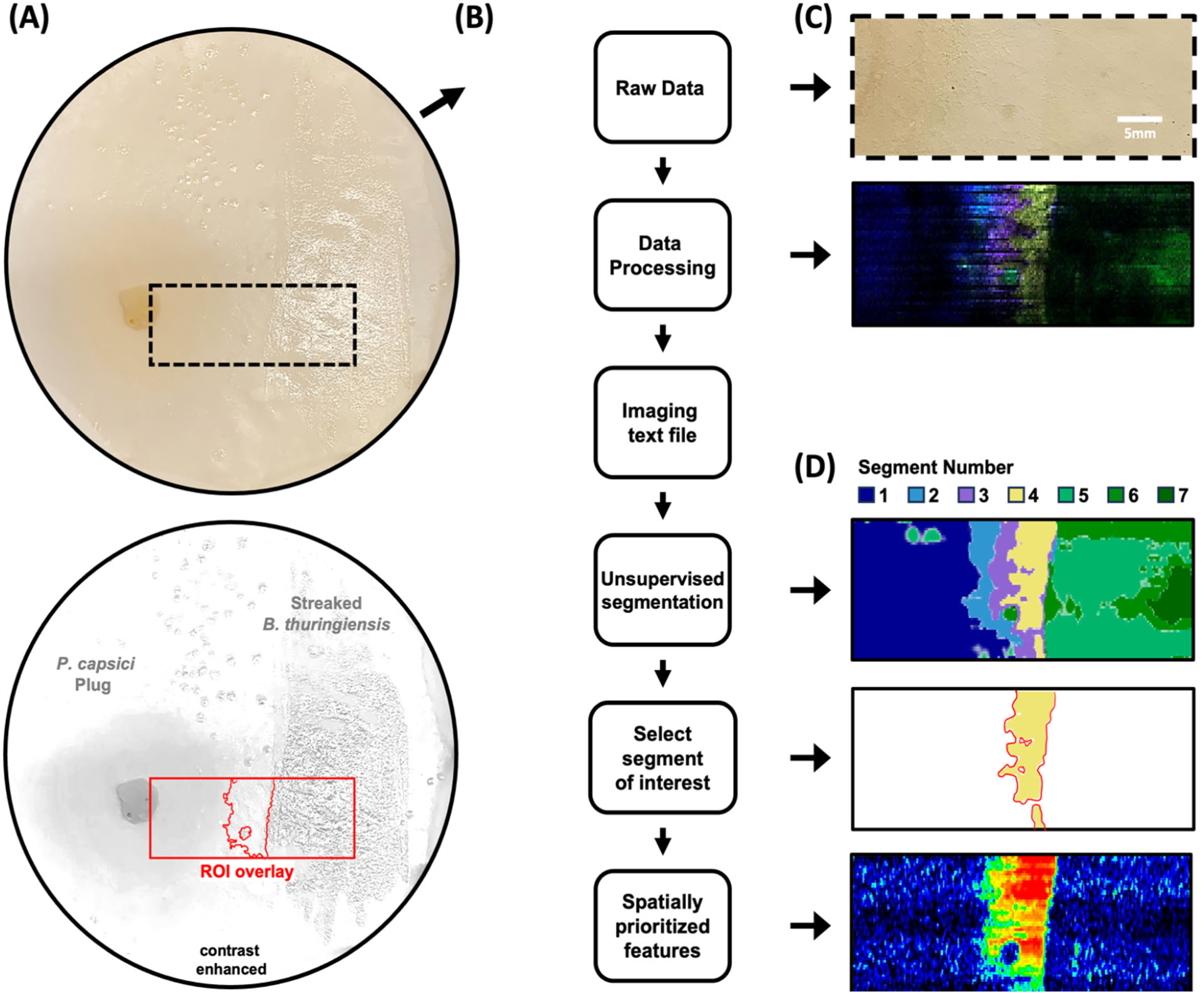
Segmentation workflow for spatiochemical profiling of *P. capsici* and *B. thuringiensis* co-culture. **(A)** Optical image (top) of co-culture grown on nylon membrane scaffold to preserve spatial location of chemical signals during sample transfer and data acquisition. Contrast enhanced image underscores the locations of the two species and the area being analyzed during DESI-MSI. **(B)** Data analysis workflow for application of the unsupervised segmentation algorithm. Raw data is processed in HDimaging before being imported into R, where the cardinal package performs an unsupervised segmentation algorithm using spatially shrunken centroids to determine regions with distinct chemical profiles. **(C)** Ion overlay displays selected ion images; different colors correspond to different molecular signals. **(D)** Output of spatial segmentation analysis, wherein 7 distinct spatiochemical regions were determined in an unbiased manner by the algorithm. Scale bars are as shown.

### Distinct metabolic profiles identified within pathogen/BCA co-culture

From the 4000 total analyte features measured from broadscale DESI-MSI analysis, 584 spatially-relevant analyte signals associated with the metabolic states of both *B. thuringiensis* and *P. capsici* were identified via unsupervised segmentation. The location-resolved nature of these signals can be visualized in the form of ion overlays, wherein each of the seven segments occupies discrete regions within the imaged area (**Fig. 5A**). Each of the 7 segments displayed unique chemical profiles, in terms of both the classes of analytes present as well as the intensities of the signals measured (**Fig 5B**). The significant (t-statistic value > 0) analyte signals present in Segment 1 (378 features) include lipids of the lysophosphatidylcholine (LysoPC) class as well as lysophosphatidylinositol (LysoPI) lipids, saturated free fatty acids (FFAs), and unsaturated FFAs. The significant features in Segment 2 (458 features) were predominantly non-lipid species, with the remainder being unsaturated FFAs. Five distinct classes of compounds were found to be significant in Segment 3 (130 features), with phosphatidylglycerol (PG) lipids, phosphatidylethanolamine (PE) lipids, saturated FFAs, hexosylceramide (HexCer) lipids, and non-lipid species being identified as significant in this region of interspecies interaction. Saturated FFAs, hydroxy-FFAs, PG lipids, and non-lipid species were the four classes observed in Segment 4 (47 features). Compounds from six classes were identified in Segment 5 (48 features). From the most significant number of identifications to the fewest, the classes are: PG lipids, saturated FFAs, PE lipids, unsaturated FFAs, HexCer lipids, and non-lipid species. Segment 6 (37 features) is comprised of saturated FFAs, unsaturated FFAs, non-lipid species, and hydroxy-FFAs. The significant constituents of Segment 7 (58 features), occupying the region furthest from the oomycete-dominated region of Segment 1, was identified to be comprised of saturated FFAs, unsaturated FFAs, PG lipids, hydroxy-FFAs, and LysoPC lipids.

**Figure 5.**
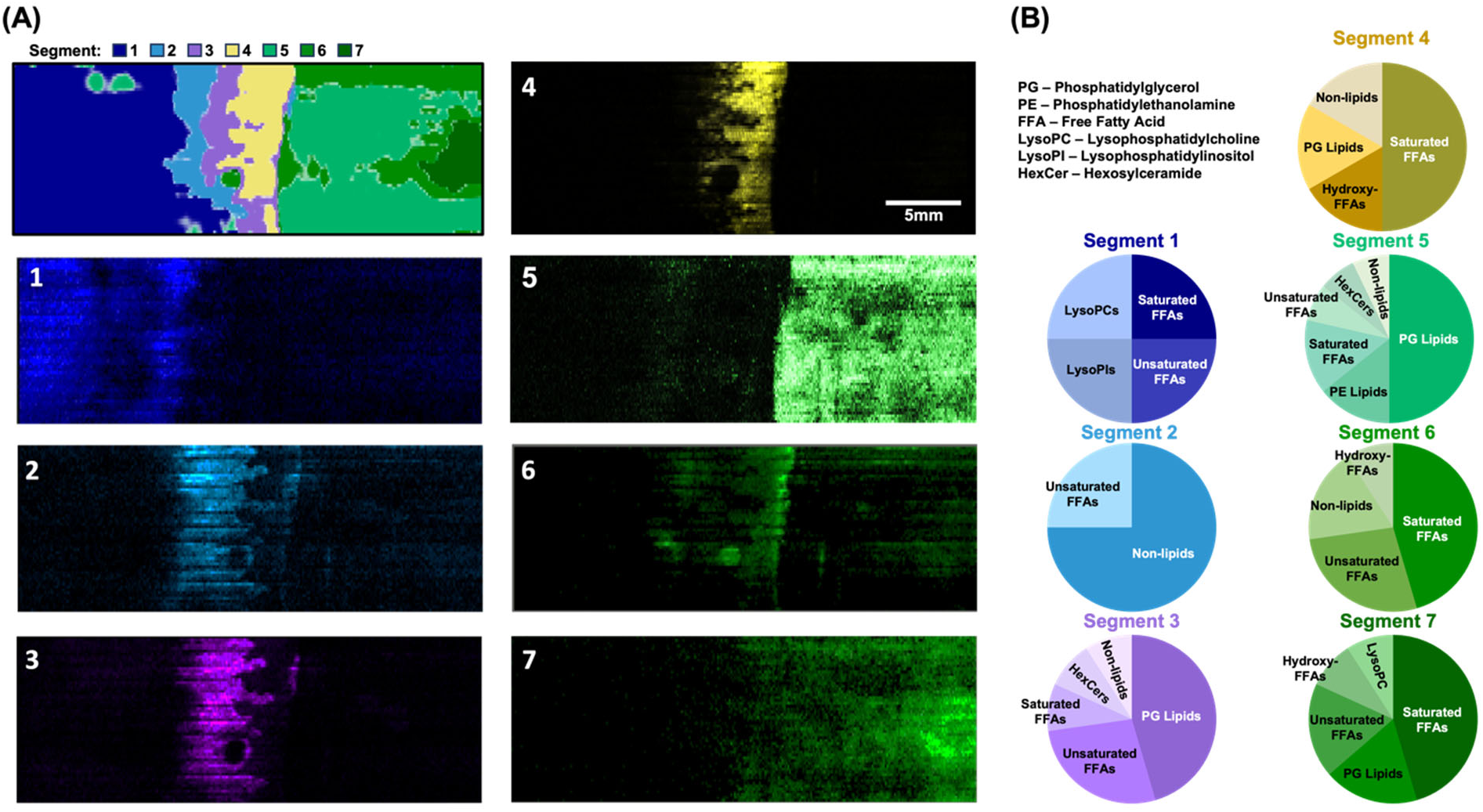
Breakdown of metabolic phenotypes identified by segmentation algorithm. **(A)** Output of segmentation algorithm (top, left) and ion overlays (1-7) for signals found in each of the seven regions determined by the SSC-based k-means clustering. Ion overlays serve to convey exact spatial localization of signals contributing to the segments identified by image analysis algorithm. **(B)** Class breakdown of identified features comprising the significant signals found within each of the phenomic regions unbiasedly determined through segmentation. Pie charts represent class breakdown for top 20 most significant features. Significance was calculated via two-sample t-test comparing the intensity of a feature within a segment area to the rest of imaged area, with identifications of signals being determined though mass accuracy calculations using a 10 ppm threshold.

For the two distal regions of the image (Segments 1 and 7), a limited number of non-lipids were observed, with all of the significant analytes identified in these two segments being lipids. Based on the optical image overlay, these regions are associated with areas principally occupied by each the two individual species, *P. capsici* and *B. thuringiensis,* respectively. In these distal regions, FFAs dominate the observed signals in terms of measured abundances, with both saturated and unsaturated FFAs as well as both even and odd chain FFAs being represented. In contrast to these observations, the regions marked by interspecies interaction (Segments 2-4), have major significant signals associated with non-lipid species. Two of the non-lipid species found to be significant within Segment 2 include signals associated with hexose and heptose sugars. These sugars are identified primarily by accurate mass measurements, and thus represent numerous possible isomers which would necessitate additional experiments (e.g., chromatography, tandem MS, ion mobility) to inform the specific sugar isomers present. By comparing the chemical class distribution associated with each segment, information surrounding the implications of altered metabolic states brought about by plating these microbial species together can be discerned. One specific chemical example for PG lipids is discussed in further detail in the next section.

### Heterogeneous lipid distribution revealed for phosphatidylglycerol class

Spatial distributions of individual *m/z* values and different analytes can be visualized as heat maps which reveals analyte specific locations within the imaged area (**Fig. 6**). One of the major lipid classes identified within the co-culture by DESI-MSI were PG lipids. Twelve PG lipids representing six different chain lengths with three discrete degrees of unsaturation were found to localize in specific regions within the co-culture. This heterogeneous lipid distribution implies varying functionality resulting from structural differences and serves to underscore the statistical observations calculated during image analysis. The saturated PG lipids PG 29:0, PG 30:0, and PG 31:0, as well as the unsaturated lipids PG 32:2, PG 33:2, and PG 34:2, were all primarily concentrated in the area of interspecies interaction located between the BCA and the oomycete (Segments 2, 3, and 4). By contrast, the saturated PG lipids (PG 32:0, PG 33:0 and PG 34:0) were all principally found within the three regions qualitatively associated with *B. thuringiensis* (Segments 5, 6, and 7), whereas the three monounsaturated lipids identified were co-distributed in the regions associated with both the oomycete and the areas of interspecies interaction (Segments 1, 3, and 4). Additional representative heat maps for FFA lipids can be found in **Supplementary Fig. S2**.

**Figure 6.**
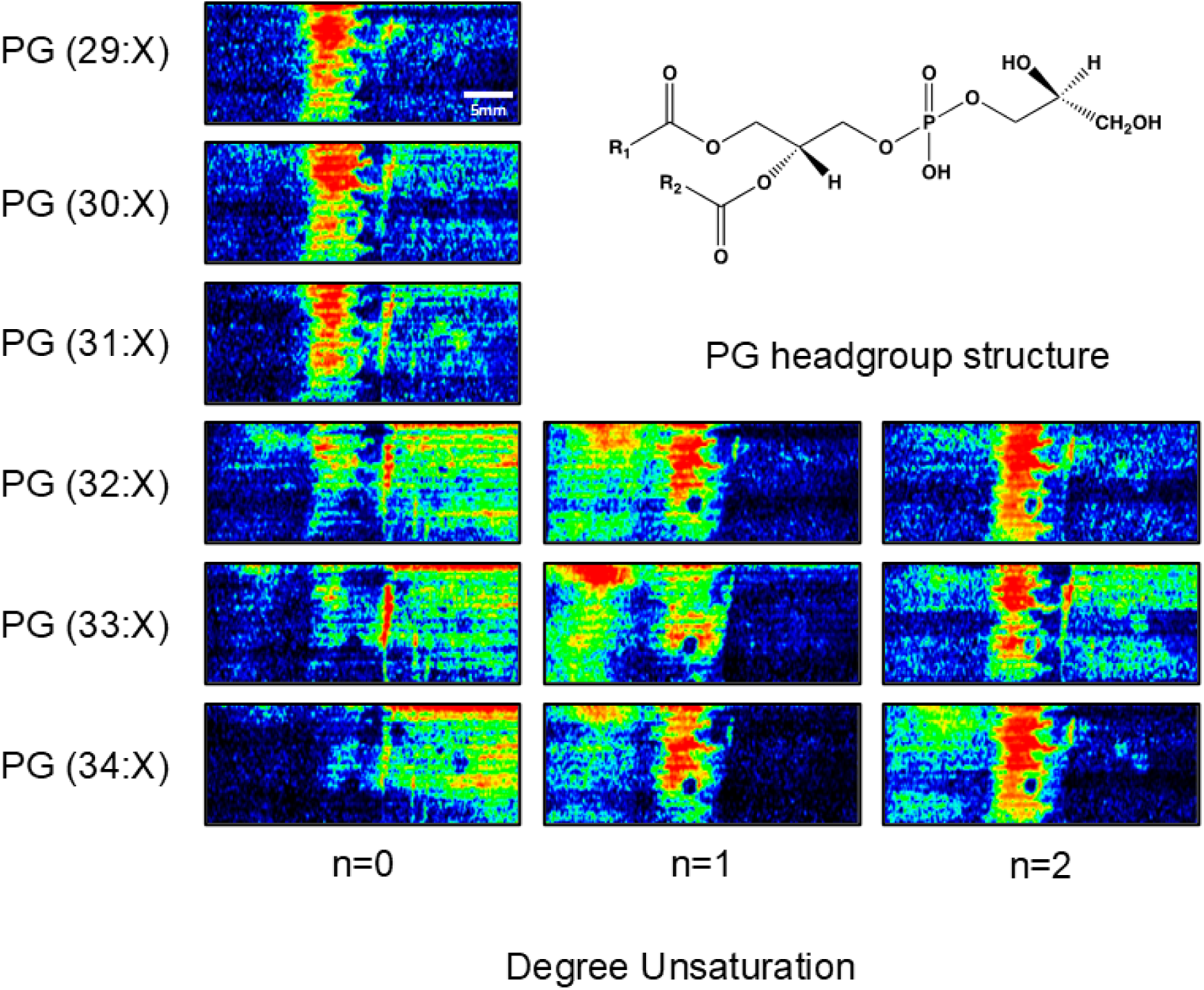
Heat maps of phosphatidylglycerol lipids reveal heterogeneous distribution. Location-resolved heat maps are produced through spatial normalization to an internal standard present in the DESI spray solution, allowing for heat maps to account for potential spray and sample inconsistencies that could occur during the course of the acquisition. Phosphatidylglycerol heat maps are organized by chain length (rows) and degree of unsaturation (columns) such that chain length increased from top to bottom and degree of unsaturation, or the sum number of double bonds present in the fatty acid tails, increases from left to right.

## Discussion

### Bacillus endophytes are potential prophylactics toward *P. capsici* infections

There is an increasing interest in suppressing *P. capsici* on tomatoes and other solanaceous species and cucurbits with rhizobacteria. Previous studies have shown that isolates of *B. amyloliquefaciens* significantly promoted the growth of seedlings and reduced pathogen load on roots, suggesting broad spectrum antifungal biocontrol traits (Syed-Ab-Rahman et al. 2019). *B. thuringiensis* is commonly used as a biocontrol agent to combat insect pests. However, new studies have revealed its potential as a preventive bacterial biocontrol agent for controlling *P. capsici* infections in tomato plants based on its antimicrobial compound production, competition with the pathogen for nutrients and space in the rhizosphere, and induction of tomatoes systemic resistance (Syed-Ab-Rahman et al. 2019; Amin et al. 2015; Martínez-Zavala et al. 2020). The presence of biological control bacteria leads to a substantial decrease in the biomass of *P. capsici* mycelium and induces significant morphological changes, culminating in the collapse of hyphae. These results suggest that biological control bacteria have a potent inhibitory effect on *P. capsici* growth and structural integrity, highlighting their potential as effective agents for managing this plant pathogen.

### DESI-MSI provides a platform for chemical analysis of interspecies bacterial interactions

In order to accurately characterize the spatial dynamics of the *B. thuringiensis* and *P. capsici* co-culture, a DESI-MSI technique was implemented for the analysis of solid-phase bacterial growth cultured on a nylon membrane. By growing the co-culture on a microporous membrane scaffold (MMS), active sample handling was minimized, allowing for the integrity of chemical signatures expressed during microbial growth to be preserved through the sample processing and analysis (Yan et al. 2017). The *in situ* molecular analysis achieved by the MMS technique combined with DESI-MSI are conducted under ambient conditions. Unlike vacuum-based MSI, the ambient nature of the DESI-MSI acquisition provides a less compromised ability to measure labile and volatile analytes, which is integral for comprehensive characterization (Ellis et al. 2019). Importantly, whereas prior work using MMS was only applied to co-cultures of gram-negative prokaryotes, this study marks the first MMS DESI-MSI investigation of an inter-domain microbial co-culture, which suggests this approach is amendable to different substrates and microbial samples (Ellis et al. 2019). This particular experimental design emphasizes the versatility of the MMS platform by allowing for the co-culturing of species with dynamic growth conditions and timescales. Here, we demonstrate the effectiveness of MMS and DESI-MSI for the metabolic probing and spatial characterization of BCA-pathogen interactions, as evidenced by the resulting spatially resolved metabolite identifications.

Due to the untargeted nature of the MMS DESI-MSI workflow developed, we were able to obtain global metabolic information for both *B. thuringiensis* and *P. capsici* through a single acquisition. However, manual curation of the resultingly large imaging datasets can be prohibitively time-consuming and can subject spatial information to undue statistical biases. To facilitate the interpretation of these large, comprehensive molecular analyses, we applied an unsupervised segmentation algorithm to deliver an unbiased assessment of the chemical ecology observed within the image, which filters the data for spatially-relevant and statistically significant features while disregarding features associated with noise. The post-acquisition image analysis algorithm provides a basis for comparison wherein features statistically enriched in particular regions result in positive values and those absent result in negative values (**Supplementary Table S1**). The MMS DESI-MSI workflow implemented in conjunction with the SSC unsupervised segmentation algorithm was found to be reproducible through inter-day biological replicates (**Supplementary Figure S3**). By isolating the most significant features within each identified region, we could prioritize the spatially relevant molecular measurements for identification.

### Spatial analysis provides insight into metabolite distributions

Our broadscale chemical imaging results provide insight into the specific chemical ecology present in the co-culture by identifying key metabolites contributing to the phenotypic profiles of each segment while simultaneously characterizing their spatial distributions. Most of the identifiable significant differences observed between the segments are due to variations in the lipid profiles, namely in terms of the lengths and the degrees of saturation of the lipid aliphatic chains, present in both the FFAs and lipid tails. For example, no significant signals identified as unsaturated FFAs were observed in Segment 4, while this class comprises a meaningful percentage of the significant signals present in every other segment. It is known that the degree of lipid saturation plays an important role in membrane fluidity, membrane permeability, and in cellular stress responses, with saturated fatty acids corresponding to more rigid and less permeable membranes that are less susceptible to oxidative stress due to the lack of double bonds along the carbon tail (Dowhan 1997). The lack of significant unsaturated FFAs present within the interspecies interaction region (Segment 4) could correspond to a cellular response occurring in the zone of inhibition favoring membranes that have the aforementioned qualities due to proximity to the pathogen. Another notable difference between the distribution of FFAs within the seven segments is the significant presence of odd-chain FFAs in every region except Segments 2 and 5. The presence of odd-chain fatty acids is to be expected based on previous investigation of the strains, though the biological interpretation of this observation within the context of spatial analysis is as yet unclear (Hamid et al. 2014; Gonzalez et al. 2022).

Without the spatiochemical information provided by the MMS DESI-MSI analysis, attributing individual signals to specific species or regions within the co-culture would be challenging. The utility of the increased depth of analysis garnered through DESI-MSI is particularly emphasized in the investigation of metabolite classes known to be produced by the two species present in the co-culture, such as PG lipids and FFAs. PG lipids are known to play an integral role in prokaryotic membrane function (e.g., charge-based self-assembly, protein translocation, initiation of replication, etc.) and there is literature precedent for the presence of PG lipids in the vegetative growth for both species studied (Hamid et al. 2014; Gonzalez et al. 2022). Our study marks the first instance where the analysis of living bacterial-oomycete co-cultures provided observation of heterogeneous spatial distributions for PG lipids and FFAs, which is directly related to the greater understanding of the biochemical processes of microbial BCAs.

## Conclusion

In this work, the effectiveness of *B. vallismortis, B. amyloliquefaciens*, *B subtilis,* and *B. thuringiensis* as growth promoting and biocontrol agents against *Phytophthora capsici* in tomato was measured. While *B. subtilis* and *B. thuringiensis* were observed to promote plant growth and provide pathogen protection against *P.capsici* in both *in vitro* and in growth chamber environments, *B. vallismortis* and *B. amyloliquefaciens* were found to be less effective in non-sterile soil greenhouse studies. The analysis of a pathogen-BCA co-culture model via DESI-MSI highlighted the interactions and chemical signaling occurring between *P. capsici* and *B. thuringiensis* by investigating the molecular interactions that serve as the foundation for the underlying biochemistry. DESI-MSI was demonstrated to be an effective analytical strategy in the spatial analysis of the bioactive compounds generated by BCAs, providing deeper insight into the benefits of BCA application to tomato cultivation.

## Materials and Methods

### BCA and pathogen culturing

All bacterial strains used in this study are listed in **Supplementary Table 1**. Single colonies of endophytic *Bacillus* isolates were routinely cultured in Luria-Bertani (LB) broth and incubated at 30 °C for 16-20 hrs with aeration at 200 rpm. Cultures were adjusted to an optical density at 600 nm 1.0 (which corresponds to approximately 10^8^ CFU / mL). Cultures were subsequently streaked on LB agar plates and incubated at 30 °C for 16 hrs for short-term storage or used directly for seed treatment.

*P. capsici* was grown from 5 mm plugs (stock stored in distilled sterile water at room temp) on V8 juice agar (20% V8 juice, 0.2% CaCO_3_, 1.5% Agar) at room temperature in the dark for 5-7 days. Zoospore suspensions were created by saturating the plates with cold sterile distilled water and carefully extracting the mycelial surface. The concentration was adjusted to 10^4^ zoospores CFU/mL. Dual culture studies used a 5 mm V8 agar plug of actively growing *P. capsici* placed at the center half of a 70 mm Petri dish containing a 50/50 mix of LB/V8.

### Evaluation of biocontrol bacterial isolates antagonism of *Phytophthora capsici*

To test the effects of BCA on plant growth and protection against *P. capsici* pathogen, tomato seeds (*Solanum lycopersicum* L. “Rutgers”) were surface sterilized in 10% sodium hypochlorite for 10 mins and rinsed three times for 60 s in double distilled water (ddH_2_O) until the smell of bleach was no longer present. Seeds were blotted dry using sterilized paper towels and placed in 50 mL conical tubes containing 30 mL of selected BCA bacterial suspension of 10^8^ for 24 hrs at 4 °C. Control treatments comprised of tomato seeds soaked in sterile LB broth with no bacteria. Seeds were air dried in a sterile laminar hood for 2 hrs prior to being sown in heat-sterilized Miracle-Gro® potting mix in 9 cm containers with a replication of seven individual plants per treatment and arranged in a completely randomized design. Studies were carried out in both growth chambers set at 26 ± 3 °C using heat-sterilized soil and in greenhouse environment using non-sterile soil.

In order to produce pathogen-infested soil, plants were drenched at the soil line with a liquid suspension of *P. capsici* zoospores at a concentration of 10^4^ spores per mL harvested from actively growing *P. capsici* mycelia. Plants were BCA treated as explained above, maintained in a growth chamber for six weeks, and then moved to the greenhouse environment, where they were maintained for an additional four weeks (ten weeks total), after which plant biomass was measured by root weight, shoot weight, and total dry weight.

Plant growth-promoting activity and pathogen protection were evaluated on Murashige and Skoog (MS) agar (MS basal media; Sigma-Aldrich, 0.03 % phytagel, pH 5.7) culture plates (11×11 cm square). Two 5 mm plugs of actively growing *P. capsici* were placed equal distances apart approximately 35 mm from the bottom of the plate. Plant growth was measured by root and shoot height of 7 day and 14 day-old seedlings using Image J software which was used to convert pixels to millimeters in optical images.

### Dual culture *in-vitro* visual analysis

A 5 mm agar plug containing actively growing *P. capsici* mycelium was positioned at the midpoint of a 50/50 V8/LB agar plate. It was given 48 hrs to grow before streaking a 20 mL culture of endophytic bacteria roughly 3 cm away from the edge of the mycelial plug. The plates were placed in a dark environment at an ambient temperature and monitored for 72 hrs. A dissecting microscope (Olympus XZS-ILLD2) coupled with an HD 4K camera was used to measure and analyze the inhibition zone and assess antagonistic activity between the BCAs and *P. capsici*.

### DESI-MSI Sample Preparation and Acquisition Settings

To prepare co-culture for DESI-MSI analysis, a 5 mm plug of *P. capsici* was placed on top of a nylon membrane set atop a 50/50: LB:V8 agar base. Subsequently, the same membrane was streaked with *B. thuringiensis*, and the co-culture was allowed to incubate at room temperature in low light conditions for five days. Nylon membranes used in this work had a porosity of 0.45 μM and were 47 mm in diameter (Sigma-Aldrich, Z290807). Prior to imaging, the membrane and co-culture were removed from agar and cut to the desired size using mayo scissors. Once cut, the membrane and bacteria were allowed to dry for 5 min and then adhered to a glass microscope slide via double-sided Scotch® tape. The mounted membrane was then imaged directly.

Imaging experiments were carried out on a Synapt G2-S High Definition Mass Spectrometer (Waters Corporation) fitted with a prototype DESI-MSI probe (Tillner et al. 2017). Optimized DESI source parameters were found to be a capillary voltage of −5 kV, 150 °C source temperature, 0.45 bar of N_2_ gas flow, and an ablation diameter of 0.7 mm. DESI geometry was set to an x,y,z positional orientation of −2,+2,+2.75, respectively, and a sprayer angle of 70°. The DESI ionization solvent consisted of 90/10 acetonitrile/water, 0.2 ng/μL leucine-enkephalin, and 0.1% ammonium hydroxide. Pixel dimensions of 100 μm x 200 μm were used and with a raster rate of 150 μm/sec being implemented. The instrument was mass calibrated with sodium formate salt clusters to a 95% confidence band and RMS residual mass of <0.5 ppm. All acquisitions were carried out in the mass range of *m/z* 50-1200, with experiments being performed using negative polarity.

### Data Processing and Spatiochemical Analysis

All raw imaging files were processed using HDImaging software (Waters Corporation), with the 4000 most abundant features present within the MSI being investigated. Lock mass adjustments were performed using leucine-enkephalin internal standard. All subsequent spatial analysis on processed files was performed in R studio (RStudioTeam 2024, v1.1.383). Feature intensities were normalized to the total ion chromatogram and then normalized to the leucine-enkephalin internal standard to account for sprayer inconsistencies and variation in sample morphology. Features were peak picked using a 10 ppm window. Spatially normalized heat maps were produced by plotting the intensity associated with peak picked features for each pixel location. Prior to segmentation, the initial number of families (k) and the shrinkage parameter (s) were determined empirically for experiments (Bemis et al. 2015). Unsupervised segmentation was achieved using spatial shrunken centroids analysis via the Cardinal MSI package, allowing for the identification of distinct spatiochemical regions within the co-culture (Bemis et al. 2015). The significance of the signals present within each region was determined by a two-sample t-test comparing the intensity of a feature within a segment area to the rest of the MSI. Tentative identifications for features were generated by searching measured *m/z* values against the Kyoto Encyclopedia of Genes and Genomes, METLIN, LipidMaps, and ChemSpider to determine tentative identifications. A threshold of 10 ppm mass accuracy and 80% isotopic similarity were used for all tentative identifications.

## Acknowledgments

This work was supported in part using the resources of the Center for Innovative Technology at Vanderbilt University. Collaborative support was provided by Waters Corporation as a Waters Centre of Innovation.

## Author Contribution

J.R. and H.S. conceptualized the study. J.R., H.S., M.M., J.C.M., and J.A.M. designed the research, planned experiments, and interpreted the results. J.R. and H.S. conducted major experiments and data analysis, with contributions from D.A. and P.E. The manuscript was written collaboratively by all authors, all of whom have approved of the final version of the manuscript.

## Supplementary data

The following materials are available in the online version of this article.

**Supplementary Figure S1.** Normal *Phytophthora capsici* morphology on 50/50 mixed plates of LB and V8.

**Supplementary Figure S2.** Additional heatmaps of interest.

**Supplementary Figure S3.** Segmentation output of biological replicates.

**Supplementary Table S1.** List of endophytic biological control agents.

**Supplementary Table S2.** Feature List and Statistics.

## Funding

Financial support for aspects of this work was provided by Department of Energy (DOE) awards DE-SC0019404 and DE-SC0022207 and National Institute of Food and Agriculture (NIFA) award 2021-38821-34595.

## Data availability

The data underlying this article are available in the article and in its online supplementary material.

